# Virtual reality-based sensorimotor adaptation shapes subsequent spontaneous and naturalistic stimulus-driven brain activity

**DOI:** 10.1101/2021.12.13.471903

**Authors:** Meytal Wilf, Celine Dupuis, Davide Nardo, Diana Huber, Sibilla Sander, Joud Al-Kaar, Meriem Haroud, Henri Perrin, Eleonora Fornari, Sonia Crottaz-Herbette, Andrea Serino

## Abstract

Our everyday life summons numerous novel sensorimotor experiences, to which our brain needs to adapt in order to function properly. However, tracking plasticity of naturalistic behaviour and associated brain modulations is challenging. Here we tackled this question implementing a prism adaptation training in virtual reality (VRPA) in combination with functional neuroimaging. Three groups of healthy participants (N=45) underwent VRPA (with a spatial shift either to the left/right side, or with no shift), and performed fMRI sessions before and after training. To capture modulations in free-flowing, task-free brain activity, the fMRI sessions included resting state and free viewing of naturalistic videos. We found significant decreases in spontaneous functional connectivity between large-scale cortical networks – namely attentional and default mode/fronto-parietal networks - only for adaptation groups. Additionally, VRPA was found to bias visual representations of naturalistic videos, as following rightward adaptation, we found upregulation of visual response in an area in the parieto-occipital sulcus (POS) in the right hemisphere. Notably, the extent of POS upregulation correlated with the size of the VRPA induced after-effect measured in behavioural tests. This study demonstrates that a brief VRPA exposure is able to change large-scale cortical connectivity and correspondingly bias the representation of naturalistic sensory inputs.

**Significance statement:** In the current work, we tested how a brief sensorimotor experience changes subsequent brain activity and connectivity. Using virtual reality (VR) as a tool for sensorimotor training opens a window for creating otherwise impossible sensory experiences and sensorimotor interactions. Specifically, we studied how VR adaptation training in ecological conditions modulates spontaneous functional connectivity and brain representation of naturalistic real-life-like stimuli. Previous adaptation studies used artificial, lab-designed setups both during adaptation and while measuring subsequent aftereffects. Testing brain response while observing naturalistic stimuli and in resting state allowed us to stay as close as possible to naturalistic real-life-like conditions, not confounded by performance during a task. The current work demonstrates how rapid changes in free-flowing brain activity and connectivity occur following short-term VR visuomotor adaptation training in healthy individuals. Moreover, we found a link between sensory responses to naturalistic stimuli and adaptation-induced behavioural aftereffect, thus demonstrating a common source of training-induced spatial recalibration, which affects both behaviour and brain representations of naturalistic stimuli. These findings might have meaningful implications both for understanding the mechanisms underlying visuomotor plasticity in healthy individuals and for using VR adaptation training as a tool for rehabilitating brain-damaged patients suffering from deficits in spatial representation.

## Introduction

“No man ever steps in the same river twice, for it is not the same river and he is not the same man” says the ancient Greek statement attributed to Heraclitus. Indeed, our every-day sensory experiences and interactions with the world shape the way we act and perceive. How do these interactions forge our brain? Neuroimaging studies targeted changes in brain activity and connectivity following novel sensory experience. There is ample evidence for experience-induced modulations on resting state connectivity. For example, visual perceptual learning was shown to modulate spontaneous connectivity between the visual and fronto-parietal networks engaged by the task (Lewis *et al*., 2009). Associative cortical areas and hippocampus increased connectivity following a visual encoding task (both in real life Tambini *et al*., 2010; and in virtual reality Gauthier *et al*., 2020). Frontoparietal and cerebellar networks functional connectivity strengthened after exposure to a visuomotor adaptation (Albert *et al*., 2009). In addition to changes in resting-state connectivity, sensorimotor training was shown to modulate subsequent task-induced activations. For instance, activation in motor regions was upregulated following motor sequence learning (area M1 in Karni *et al*., 1995; and premotor cortex in Berlot *et al*., 2020). Visual motion aftereffect was found in area MT following adaptation to a moving stimulus (Tootell *et al*., 1995), and auditory frequency discrimination training was shown to enhance auditory activation in proportion to performance gain (Jäncke *et al*., 2001). However, previous studies on experience-induced modulations focused mainly on local effects, restricted to the specific task and brain region being trained. It remains unknown how recent sensory experience affects subsequent representation of task-free naturalistic stimuli, possibly implemented across long-range connections.

Sensorimotor training is not the sole responsible of behavioural and neural changes. Some neurological pathologies lead to biased sensory representations and aberrant interactions with the environment, thus entailing continuously altered sensory experience. A paradigmatic case is hemispatial neglect syndrome (‘neglect’). In this unique pathology, right hemisphere damage causes patients an inability to attend to and interact with stimuli in the left side of space. In many cases, the lesion affects key fronto-parietal regions in the attentional networks (Corbetta & Shulman, 2011), while keeping the sensory cortices intact. Nevertheless, the effects of the lesion span well beyond the focal damage, causing a general imbalance of brain activity and connectivity (Baldassarre *et al*., 2014; Lunven & Bartolomeo, 2017; Xu *et al*., 2019). More recent accounts discovered an imbalance between brain networks in neglect patients at rest, as manifested in reduced anticorrelation between attentional networks and the default mode network (DMN; Baldassarre *et al*., 2014; Siegel *et al*., 2016). These recent findings emphasize the role of fronto-parietal attentional and DMN regions in mediating representation and processing of multisensory inputs for well-functioning sensorimotor interactions.

In the current study, we implement a novel virtual reality-based visuomotor adaptation training to affect the way healthy people interact with and represent the environment. We ask whether and how such training, which induces a shift of reference frames, modifies large scale brain networks connectivity and the processing of naturalistic stimuli. To this aim, we base on a prominent method for studying visuomotor plasticity in healthy individuals, as well as for rehabilitating neglect patients, called prism adaptation (‘PA’). It consists in performing repetitive goal-directed movements while wearing prismatic lenses that induce a lateral shift of visual inputs by a fixed degree. To compensate for this discrepancy, participants recalibrate their movements in the direction opposite the shift, resulting in spatial biases in open-loop reaching tests after prism removal (‘PA aftereffects’; cf. Redding & Wallace, 1996). PA aftereffects span well beyond the motor domain, as it induces a shift in reference frames which affect various attentional and perceptual tasks, in both neglect patients and in healthy individuals (Jacquin-Courtois *et al*., 2013; Michel, 2015), and persist long after the adaptation training session had ended (> 40 minutes) (Schintu *et al*., 2014). Previous studies show that the aftereffects of PA manifest also in brain activity as changes in activation of the inferior-parietal lobule (IPL; Crottaz-Herbette *et al*., 2014; Crottaz-Herbette *et al*., 2017b), a key area at the intersection between the attentional networks and the DMN. In addition to its effect on task-induced activation, PA has been recently found to modulate task-free resting-state connectivity, inducing enhanced decoupling between the attentional networks and the DMN (Wilf *et al*., 2019) or a modulation of the right/left fronto-parietal networks (Schintu *et al*., 2019; Tsujimoto *et al*., 2019; Gudmundsson *et al*., 2020; see Panico *et al*., 2020 for review).

These previous studies largely used artificial, non-ecological setups, both during the PA training phase, and while testing for task-related aftereffects in fMRI. We propose that in order to gain better understanding of real-life-like aftereffects of sensorimotor adaptation, both the adaptation training and the fMRI paradigms testing its brain effects, should adopt a naturalistic approach (cf. recent opinion by Nastase *et al*., 2020). Thus, in order to enhance the ecological nature of the adaptation training, we here implement PA training in an immersive VR environment (VRPA) embedding dynamic visual targets and a gamified activity. In three different groups of participants, we introduced a rightward, a leftward or a sham visuomotor rotation, and assessed the related behavioural aftereffects (see *Figure 1a,b*). To unravel the neural aftereffects of VRPA, we performed fMRI sessions assessing task-free brain activity before and after the training, namely, spontaneous brain activity and brain response to free-viewing of naturalistic stimuli (*Figure 1b,c*). We investigated VRPA-induced changes in cortical functional connectivity patterns, changes in activation pattern in response to naturalistic stimuli, and their link with well-known behavioural proprioceptive adaptation aftereffects. To further examine whether these modulations are hemisphere-specific, depending on the spatial directionality of the VRPA shift, we compared the results between the rightward and leftward VRPA groups. Using this experimental design, we characterize the mechanisms underlying short-term VR experience-induced modulations in free-flowing brain activity.

**Figure 1.**
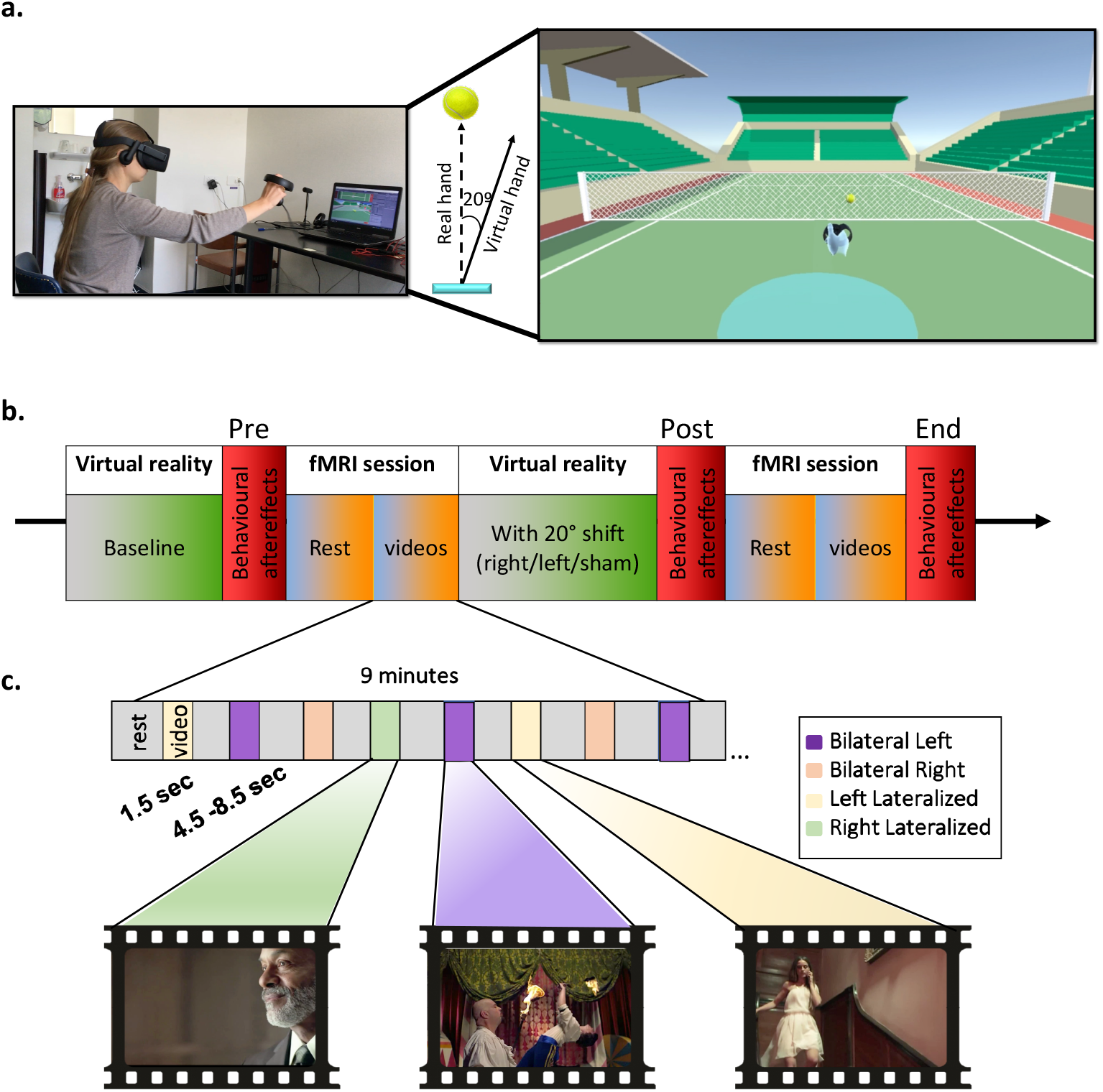
Experimental paradigm and procedure. (a) Schematic representation of virtual reality prism adaptation (VRPA) training. Participants were immersed in a 3D virtual environment and had to catch dynamic targets (tennis balls) using a virtual hand. During adaptation sessions, a 20° rightward/leftward angular shift was introduced between the participants’ real hand and the virtual hand. (b) Experimental procedure. The session always began with baseline VR training with no shift, followed by spatial bias tests including manual pointing straight ahead with eyes closed. Afterwards, participants underwent resting-state fMRI and a free viewing of a sequence of naturalistic videos. They then performed VRPA outside the scanner, with either right (n=16), Left (n=15); or no shift (n=14), followed by an identical set of behavioural tests and fMRI experiments. At the end of the session, behavioural aftereffects were again measured outside the scanner. (c) Scheme of naturalistic videos experiment. Videos lasted 1.5s and contained a salient stimulus on either the right/left side of the screen, or on both sides of the screen, with the more salient stimulus on the right/left side (20 videos in each category; videos taken from (Nardo et al., 2016; Nardo et al., 2019)).

## Results

### Behavioural VRPA aftereffects

First, we assessed the behavioural aftereffects induced by the VRPA training. Participants exhibited consistent lateral biases in manual straight ahead pointing with eyes closed following either right or left VRPA training, but not following sham VRPA, thus demonstrating a clear aftereffect due to the sensorimotor adaptation. In the leftward VRPA group, aftereffect manifested in systematic rightward deviation in pointing straight ahead movements (*Figure 2a*; main effect of experimental phase: F(2,28) = 16.4; p < 0.001), while the rightward VRPA group presented leftward deviations following adaptation (*Figure 2b*; main effect of experimental phase: F(2,30) = 13.5; p < 0.001). No systematic pointing error was induced by the Sham VRPA (*Figure 2c*; F(2,26) = 1.9; P > 0.15). Importantly, a residual pointing bias was still evident in both adaptation groups even at the end of the experimental session, i.e., approximately 40 minutes after VRPA (*Figure 2a,b*; green; in line with (Schintu *et al*., 2014)). These behavioural results validate that the spatial recalibration induced by the VRPA training was effective when participants were undergoing subsequent fMRI experiments and that some of the effect remained throughout the entire ‘post’ fMRI session.

**Figure 2.**
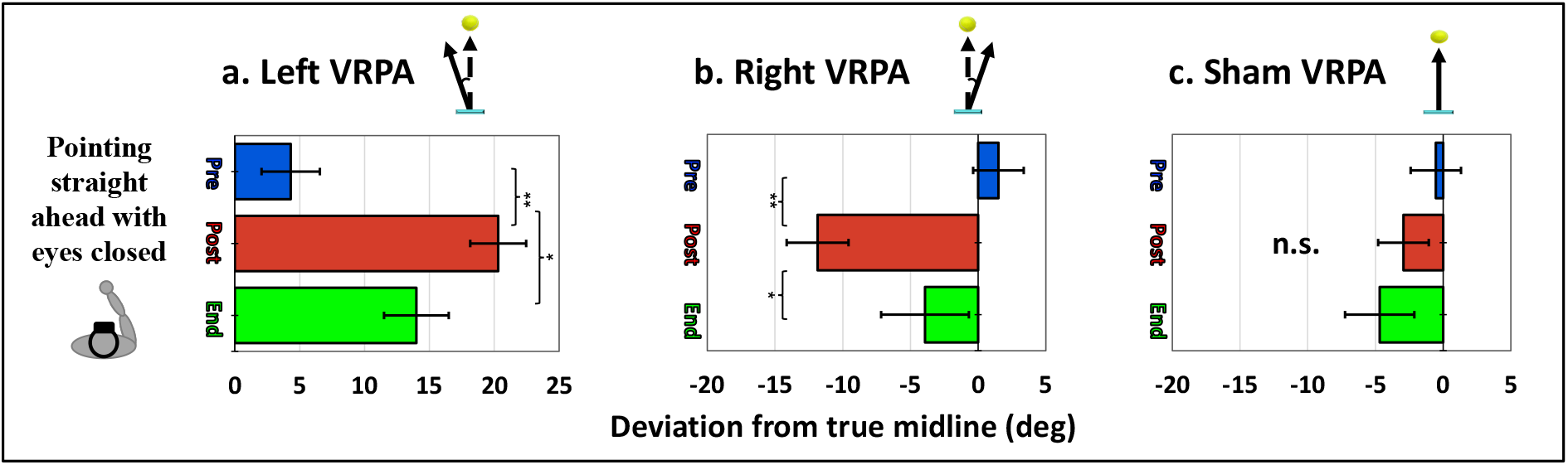
behavioural aftereffects following rightward, leftward, and sham VRPA. Participants had to point straight ahead with eyes closed on three timepoints during the experimental session: after baseline VR practice (‘Pre’), after VRPA training (‘Post’), and at the end of the experiment after the fMRI session (‘End’, ~40 minutes following adaptation). Bar graphs represent average pointing errors in each group (a) left VRPA (N=15), (b) right VRPA (N=16), (c) sham VRPA (N=14). Positive values denote rightward deviations, and negative values denote leftwards deviation. * p < 0.05; ** p < 0.001.

### VRPA effects on resting-state connectivity

After we established the presence of behavioural aftereffects, we aimed to unravel VRPA-induced modulations in spontaneous, resting-state connectivity. To obtain a complete, unbiased, view of cortical connectivity, we represented the resting-state connectivity pattern as a matrix of correlations between a set of 48 atlas-based cortical regions of interest (ROIs), homologous between the left and right hemispheres, which were sorted according to seven primary functional networks (see *Methods* and *Figure 3a*). All three groups showed typical resting-state connectivity patterns at baseline, prior to VRPA or sham training (*Figure 3b*). However, when assessing the modulations in connectivity between pre- and post-VRPA, there was a substantial difference between the adaptation groups and the sham group: the left and right VRPA groups showed widespread decreases in connectivity (*Figure 3c; left+middle*), while the sham group showed only slight increases in connectivity, mainly between visual and motor ROIs (*Figure 3c; right*). A qualitative inspection of the modulation matrices revealed that the connectivity decrease in the VRPA groups was most prominent in the connections between the default mode network / fronto-parietal networks regions (DMN/FPN), and the two main attentional networks – the dorsal and ventral attentional networks (DAN/VAN), including the IFG and STS regions (*Figure 3c; left+middle; black square outlines*). Interestingly, while following right-VRPA the most pronounced connectivity decreases were evident within the left hemisphere and between left and right hemispheres (*Figure 3c; middle*), following left-VRPA the effect was more evident in the right hemisphere and between hemispheres (*Figure 3c; left*). Together, these results suggest that VRPA (but not sham VRPA) increases decoupling between DMN/FPN and attentional networks, with more pronounced modulation between hemispheres and within the hemisphere corresponding to the direction of the behavioural aftereffect (i.e., right hemisphere for left VRPA and left hemisphere for right VRPA).

**Figure 3:**
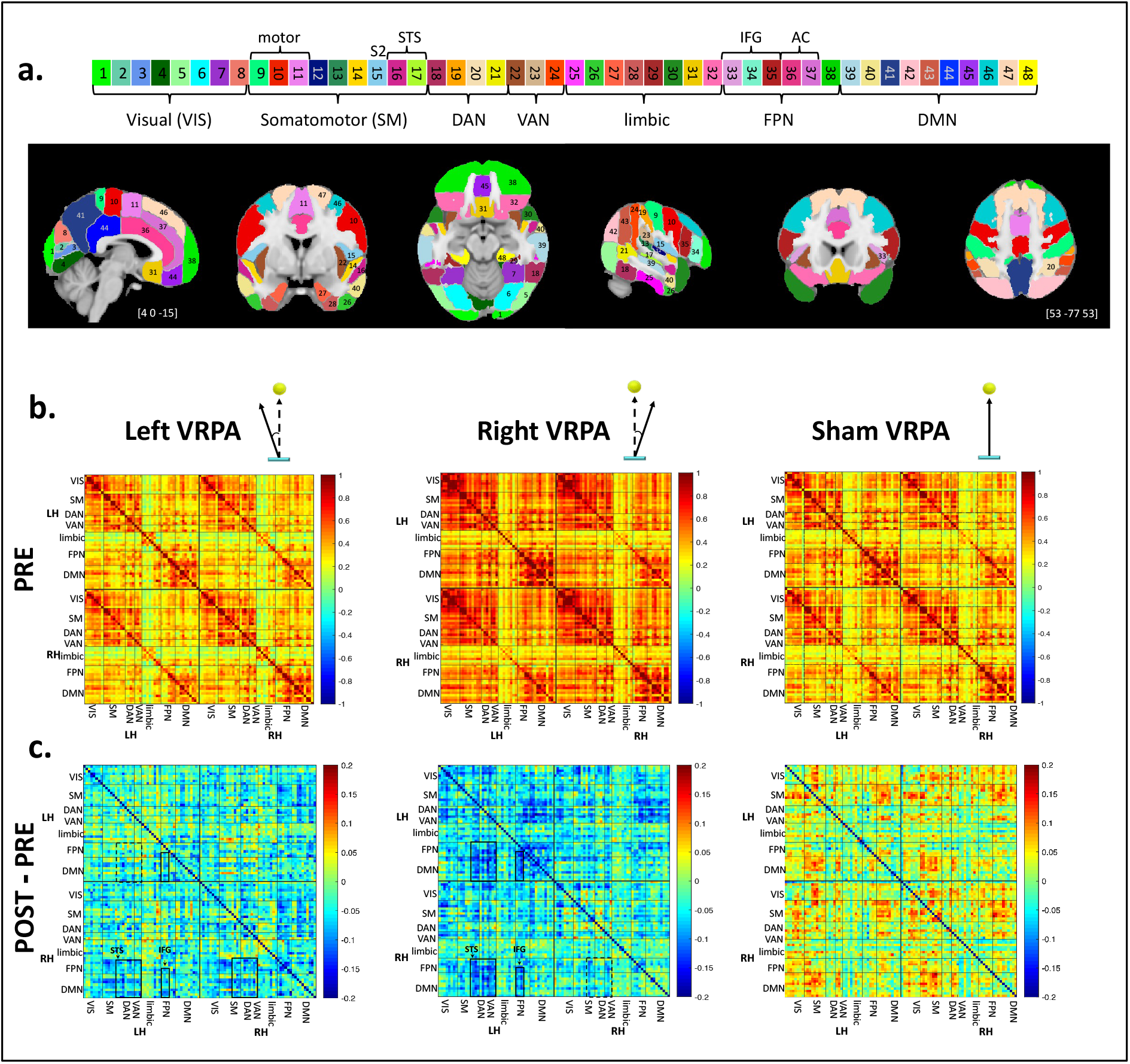
Resting-state connectivity modulations following VRPA. (a) Atlas-based cortical segmentation of MNI brain with network labelling. Color-coded Harvard-Oxford atlas regions are symmetrical across the two hemispheres. The atlas regions are labelled and principally sorted according to the seven primary functional connectivity networks suggested in Yeo et al., 2011, and within each network according to functional subsystems (see methods). (b) Mean baseline (pre-VRPA) pairwise functional connectivity matrices between atlas regions in the left and right hemispheres (N=14 for right/sham groups; N=15 for left group). Values are sorted within each hemisphere as denoted in panel a, with identical sorting for left and right hemispheres. Each quadrant represents either within hemisphere connections (LH-LH/RH-RH), or inter-hemisphere connections (RH-LH). (c) Mean pairwise connectivity modulations following VRPA (either left/right/sham VRPA groups). Blue colours denote decrease in correlations and warm colours denote increase in correlations. Note that following either type of VRPA (but not sham), there was an overall decrease in connectivity. Black squares highlight clusters of connectivity modulations between DMN/FPN and attentional networks. Homologous connections were modulated across quadrants in the left/right VRPA groups, but with different hemispheric dominance. VIS = visual network; SM = sensorimotor network; DAN = dorsal attention network; VAN = ventral attention network; FPN = frontoparietal network; DMN = default mode network; IFG = inferior frontal gyrus; AC = anterior cingulate; STS = superior temporal sulcus; LH = left hemisphere; RH = right hemisphere.

In order to quantify the VRPA-induced modulations in resting-state connectivity, we fitted a Gaussian Mixture Model to the distribution of correlation values in each connectivity matrix (see *Methods*). First, to compare the overall changes in connectivity, the means (μ values) of the fitted gaussians were compared between ‘pre’ and ‘post’ matrices in all three experimental groups (see *Figure 4a* for two single-subject examples of gaussian fitting and mean value extraction). A 2-way mixed-effects ANOVA with factors group (left/right/sham VRPA) and phase (pre/post) revealed a main effect of group (F(2,40) = 4.54; p = 0.017) and an interaction between group and phase (F(2,40) = 5.54; p = 0.007). These results demonstrate that the pattern of connectivity before and after VRPA varied across the groups.

**Figure 4.**
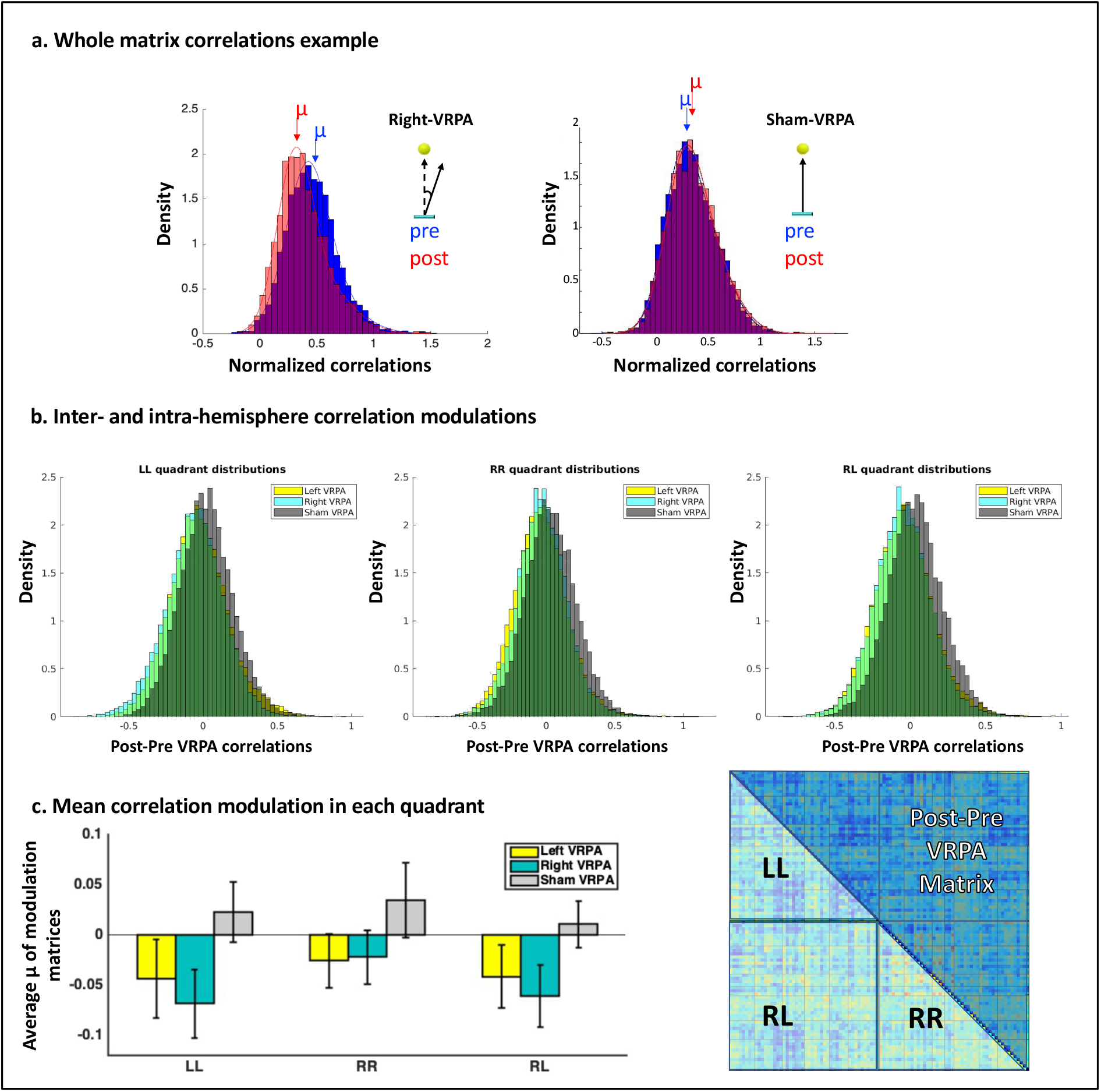
Gaussian Mixture Model results for comparing between correlation matrices. (a) Single participants example (from the right-VRPA and the sham-VRPA groups), showing the distribution of correlation values in the connectivity matrix (Fisher’s Z normalized) pre- and post-VRPA. A mixture of 2 Gaussians was fitted to each distribution, from which a μ value representing the mean was extracted and used for further statistical analysis. (b) Connectivity modulations across hemispheres. Distribution of mean modulation matrix values (post-pre VRPA) in the three groups in each of the possible connectivity types – within left hemisphere (left; LL), within right hemisphere (middle; RR), and between hemispheres (right; RL). (c) left side: average μ value of modulation matrices across participants in each quadrant for each group (errorbars denote SEM across participants). Right: a schematic example of modulation matrix with an annotation of the submatrices representing each quadrant.

Next, in order to deeply investigate connectivity changes induced by the different types of visuomotor adaptation, we zoomed in on three sub-types of connectivity modulations – within left hemisphere (‘LL’), within right hemisphere (‘RR’), and between hemispheres (‘RL’), as represented in different quadrants of the connectivity matrix (see *Figure 4c*; right). Specifically, since we aimed to quantify the size of connectivity *modulation* within each quadrant, this time we applied the gaussian mixture model analysis on *the difference matrices* instead of separately for the pre and the post phase (i.e., post-pre matrices; cf. *Figure 3c*). We first plotted the distribution of the mean modulation matrices in each quadrant (*Figure 4b*), which again revealed a differentiation in the direction of modulation between the sham and the adaptation groups, with a slightly more negative distribution for right VRPA group in the LL quadrant and for the left VRPA in the RR quadrant. To quantify potential differences between quadrants, we then performed a 2-way mixed-effects ANOVA on the individual participant modulation matrices with factors group (left/right/sham VRPA) and quadrant (LL/RR/RL) and found a main effect of quadrant (*Figure 4c*; F(2,80) = 4.6; p = 0.013). These results suggest that indeed a slightly different modulation was induced in each hemisphere and across hemispheres.

Taken together, the pattern of VRPA-induced modulations suggests an overall decrease in spontaneous resting-state connectivity, with more pronounced reductions in the hemisphere opposite to that of the shift induced by the sensorimotor training, mainly between DMN and attentional regions.

### VRPA effects on brain activity while processing naturalistic visual stimuli

After establishing VRPA effects on spontaneous activity, we tested whether the training-induced modulation of brain activity would also affect stimulus-induced representations of naturalistic stimuli, namely movie scenes. To that end, we compared the cortical responses to free-viewing of an identical series of short videos before and after VRPA training (see *Figure 1c*). The videos contained salient moving objects either on the left side, right side, or both sides of the screen. This experiment was performed only for the right-VRPA group and the sham-VRPA group (see *Experiment* 2 in the Methods). *Supplementary Figure 1* shows the pattern of activation in the right-VRPA group in response to the entire set of videos, separately for pre and post VRPA phases. As expected from naturalistic stimulation, in both pre and post phases, the activation spanned the entire hierarchy of visual processing, including parietal and frontal attentional regions (*Supplementary Figure 1*; consistent with *Nardo et al., 2016*). Crucially, an enhancement of activity emerged after the right VRPA training, specifically in the right visual cortex (*Supplementary Figure 1*; *green-black arrow*).

Indeed, a direct contrast between pre- and post-VRPA activation maps revealed a significant increase of response to naturalistic videos in an area within the parieto-occipital sulcus (POS; *Figure 5a*; MNI coordinates [20, −65, 24]). This enhancement was confined to the right hemisphere, while left hemisphere activation pattern remained similar to the pattern found prior to adaptation. Thus, visual activation following right-VRPA was biased in favor of the right hemisphere, suggesting enhanced representation of the left portion of space. We further investigated whether this modulation depended on the spatial distribution of salient stimuli in the movies (i.e., left-lateralized/right-lateralized/bilateral), or it was generalized across all video categories. When examining the different video categories separately, we found a strikingly robust effect across the video categories, with the same POS region showing enhanced activation in right-lateralized, left-lateralized, and both types of bilateral videos (*Figure 5b*). Thus, the right hemisphere visual area responded more to naturalistic stimuli following right-VRPA, regardless of whether the salient object appeared on the left or right side of the screen.

**Figure 5.**
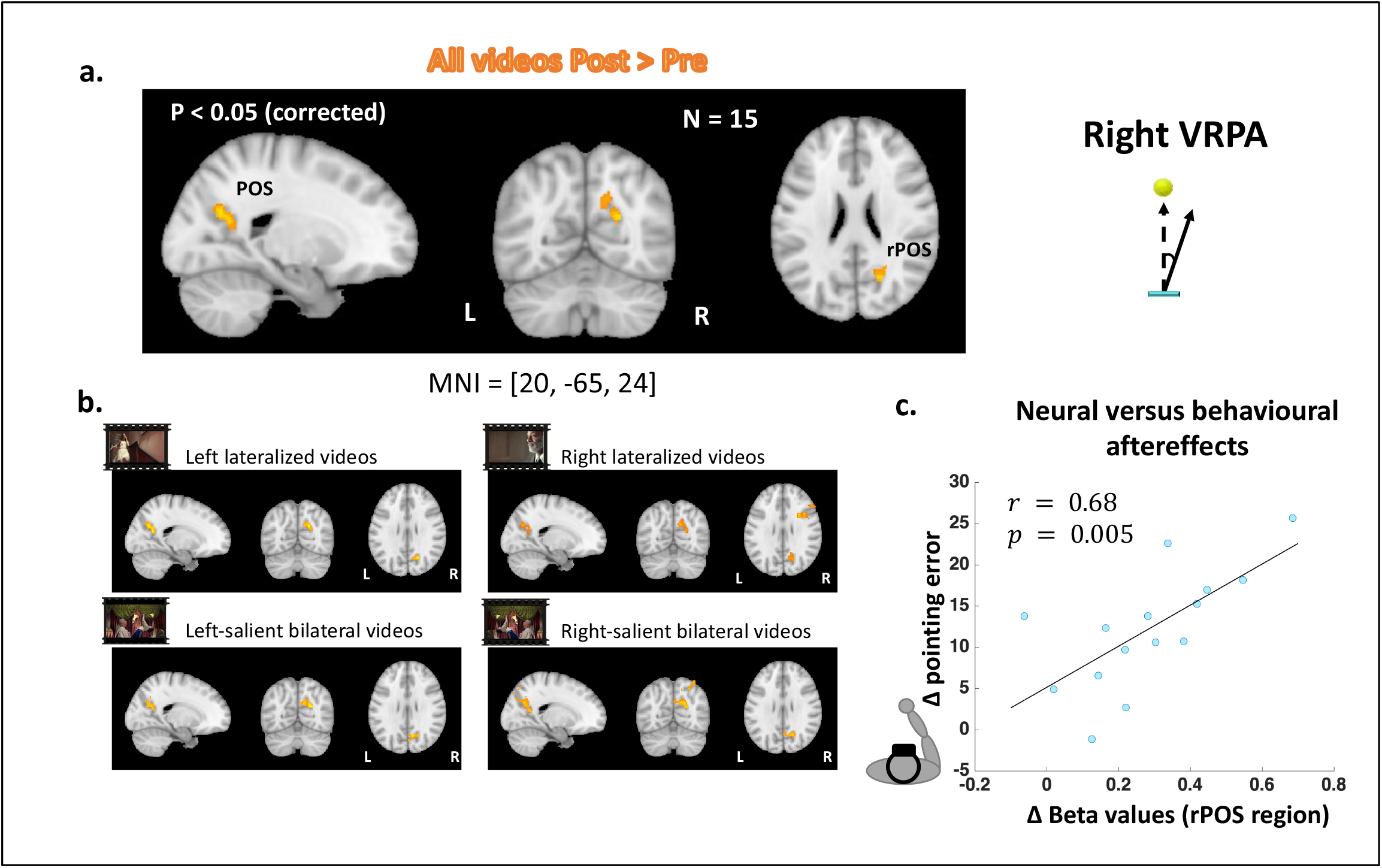
Right VRPA upregulates responses to naturalistic videos in the right POS. (a) Direct contrast between post-pre activation maps in response to all types of videos revealed enhanced activation in the right POS following right VRPA. (b) Contrast between post-pre activation maps in response to each video type. Right POS is consistently modulated. (c) Correlation between Neural and behavioural aftereffects of right VRPA. The scatter plot demonstrates the relation between (x-axis) the change in rPOS activity in response to all video types in each participant (N=15), versus (y-axis) the change in pointing error in the manual straight ahead task between pre- and post-right-VRPA (positive values here denote a more extensive leftward deviation in pointing movements in the post-session). rPOS = right Parieto-Occipital Sulcus.

Next, to probe for a link between neural and behavioural aftereffects, we compared the change in right POS activity during movie presentation (‘all videos’ condition) to the change in the pointing straight-ahead accuracy (assessing subjective body midline) between pre- and post-right-VRPA. We found a striking correlation between the extent of leftward deviation in subjective body midline induced by VRPA and the size of modulation of right POS activity while processing naturalistic visual stimuli (r = 0.68; p = 0.005; *Figure 5c*). This result provides a strong link between the effect of VRPA on sensorimotor recalibration and its effect on naturalistic visual processing.

A similar analysis was applied for the sham group to control for test-retest effects. The comparison of movie-evoked brain activity between pre- and post-sham-VRPA showed only an inconsistent enhancement in bilateral intraparietal sulcus (IPS) regions. This effect was not consistent across movie categories, i.e., it was present only for right-salient bilateral videos, and was not correlated to behavioural results (*Supplementary Figure 2*).

Finally, we further investigated how the changes in evoked activity induced by right-VRPA in the right POS during videos presentation were related to changes in its connectivity with other brain regions, using a psychophysiological interaction analysis (PPI). We found that at the ‘pre’ session (before right VRPA training), the right POS area showed no significant interactions with other areas. However, following right-VRPA, the right POS showed significant interactions with left IPL region and bilateral STS regions (*Figure 6*). These areas are part of the DMN network (*Figure 6*; green shades), which also demonstrated changes in connectivity following VRPA in the resting state connectivity analyses described above (see *Figure 3c* and *Resting State Connectivity* section). This suggests that the enhancement of POS activation in the right-VRPA group was mediated by interactions with DMN regions in the left IPL and bilateral STS. PPI analysis yielded no significant interactions in either the ‘pre’ or ‘post’ sessions in the sham-VRPA group (taking as seed region the IPS; see *Methods*).

**Figure 6.**
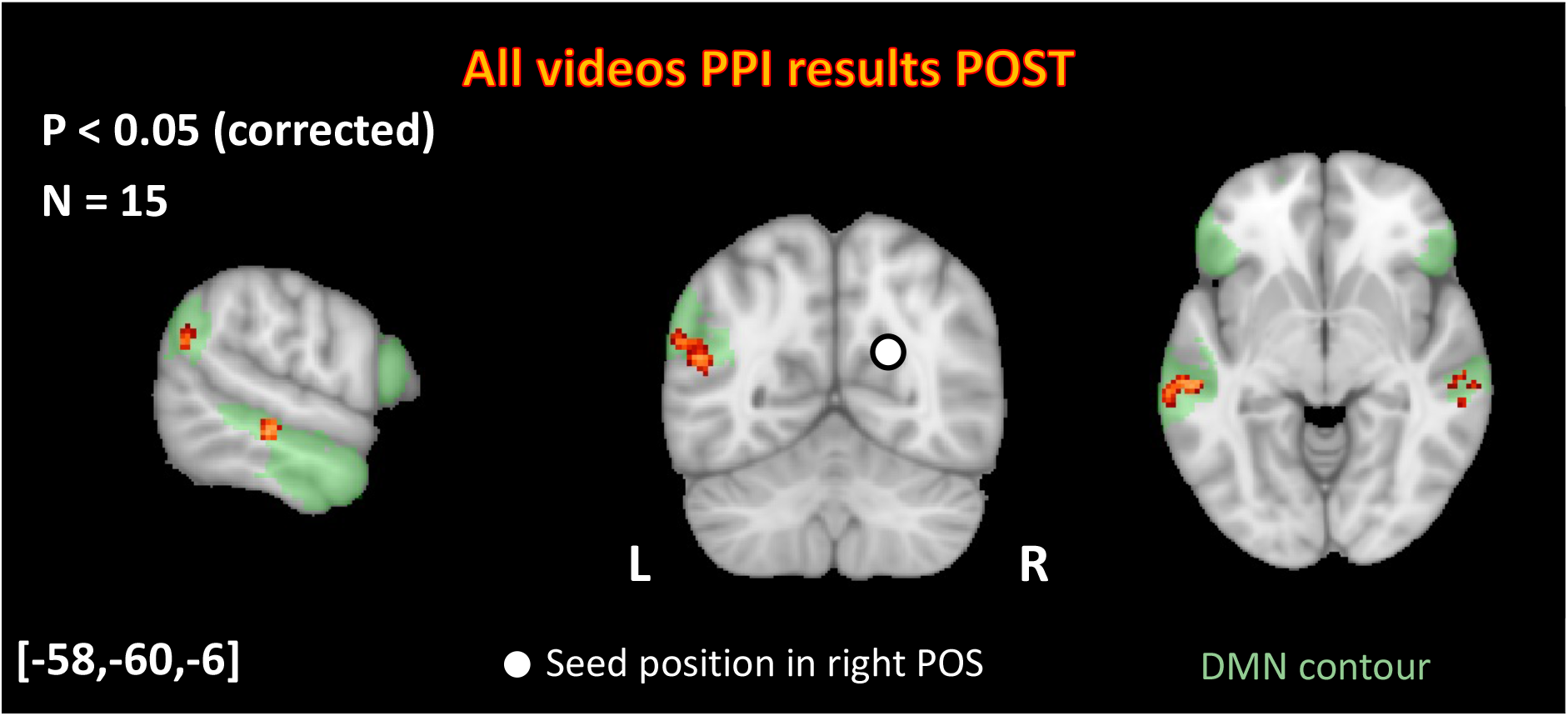
Psychophysiological Interaction analysis with right POS after right VRPA. Clusters in orange show areas that significantly interact with right POS seed (same region as in Figure 6) during video presentation following right VRPA. Areas include bilateral STS and left IPL. Light green contour shows DMN-b subnetwork as depicted in Yeo et al., (2011) 17-network parcellation (network #17). Numbers in brackets represent MNI slices coordinates.

## Discussion

VRPA successfully induced sensorimotor adaptation, as demonstrated by long lasting proprioceptive-motor aftereffects in healthy participants, that is a spatial bias in the direction contralateral to the VRPA deviation indicative of a shift in reference frame. These behavioural effects were associated with modulations in resting-state connectivity patterns, namely reduced connectivity between DMN/FPN and attentional networks, more pronouncedly in the hemisphere contralateral to the VRPA deviation. The training-induced modulations were not confined to spontaneous activity, but affected also cortical responses to free viewing of naturalistic videos. Rightward-VRPA training enhanced visual responses in a retinotopic area within the right POS, presumably reflecting enhanced representation of the left portion of space. Moreover, this hemispheric bias in visual representations was directly linked to the behavioural aftereffect, as the extent of upregulation of right POS activity during naturalistic viewing was highly correlated with the extent of leftward proprioceptive-motor biases induced by VRPA. Finally, PPI analyses showed that the right POS region interacted with DMN areas in the IPL and STS during naturalistic viewing only following right-VRPA, suggesting that changes in visual areas processing naturalistic stimuli following VRPA were mediated by these high-level, heteromodal regions. We will discuss possible interpretation and limitations of each of these results separately, and eventually suggest a model for VRPA-induced brain modulations, with potential implications for neglect rehabilitation.

### 1. Unbalancing resting state connectivity between DMN and Attentional areas

Our finding of an enhanced decoupling between DMN/FPN regions and dorsal and ventral attentional networks after VRPA (see *Figure 3b*), more pronounced in the hemisphere contralateral to the shift, is consistent with recent studies that measured standard PA aftereffects in resting state connectivity (Schintu *et al*., 2019; Tsujimoto *et al*., 2019; Wilf *et al*., 2019; Gudmundsson *et al*., 2020). We suggest that the bottom-up learning process during adaptation eventually propagates forward in the processing pathway to affect high-level, heteromodal, cortical networks. In terms of functionality, the decoupling between DMN and attention networks is crucial for execution of goal directed behaviour (Dixon *et al*., 2017; for more elaborated discussion see Wilf et al., 2019), and the extent of DMN deactivation was previously found to be correlated with learning parameters in a visuomotor adaptation task (Cassady *et al*., 2018). Additionally, STS (Luauté *et al*., 2009) and IFG (Baldauf & Desimone, 2014) regions were found to be consistently related to spatial adaptation paradigms.

Furthermore, the pattern of the hemispheric laterality visible from the connectivity modulation matrices suggests a VRPA-induced spatial recalibration and a shift in reference frame, reflecting at the brain level the mechanisms underlying PA at the behavioural level (see Redding & Wallace, 1996). This is corroborated by the fact that connectivity changes were found in networks responsible for spatial allocation of attention, such as the ventral and dorsal attentional networks (Corbetta & Shulman, 2002). This dependency on the direction of the adaptation shift is largely in line with recent findings on rightward and leftward standard PA (Schintu *et al*., 2019; Tsujimoto *et al*., 2019), who found connectivity modulations in parietofrontal attentional regions as a function of the shift direction. Findings from standard PA paradigms show also hemispheric laterality in task-induced aftereffects. For instance, Crottaz-Herbette et al., showed that right and left PA produced hemisphere-specific activation modulation in IPL region during a visual attention task (Crottaz-Herbette *et al*., 2014; Crottaz-Herbette *et al*., 2017b).

In the sham group, only slight increases in connectivity between visuomotor areas were found following training. These results are in line with the notion that sham-VRPA, without a visuomotor discrepancy, actually corresponds to an intensive multisensory-motor training in VR. Thus, the resting state patterns seem to resonate the co-activation between visual and motor areas that occur during the training (cf., Albert *et al*., 2009; Guerra-Carrillo *et al*., 2014).

In summary, exposure to VRPA modulated resting-state connectivity in high-level, heteromodal, cortical networks, in a way that reflects the spatial shift in reference frame induced by the training. In the next sections, we will propose a mechanism by which these connectivity modulations might affect processing of external sensory stimuli.

### 2. Upregulation of right POS activation during naturalistic viewing

Previous studies assessed brain activity following PA during visual attention, auditory detection and working memory tasks (Crottaz-Herbette *et al*., 2014; Tissieres *et al*., 2018). However, lab-designed task-induced activity does not correspond to the complexity of signals emerging in real life conditions (Nastase *et al*., 2020). For this reason, in the current study, we recorded brain activity that was not confound by a goal-directed task, but rather entailed implicit processing related to everyday life stimuli, as portrayed by the naturalistic videos. This approach allowed us to bridge the gap between previous results produced in highly controlled, but artificial laboratory conditions, and adaptation aftereffects in real-life-like perception, that are difficult to measure. Despite the free-flowing nature of the experience, natural viewing reliably represents sensory perception, since movie-induced brain activity is largely consistent between individuals and between movie repetitions within the same individual (Hasson *et al*., 2004; Hasson *et al*., 2010; Wilf *et al*., 2017; Strappini *et al*., 2019).

Our results highlighted the POS area as a target of VRPA-induced modulations during processing of naturalistic stimuli. This retinotopic area has a role in peripheral peripersonal space representation (Galletti *et al*., 1999), in particular during reaching or visuomotor tracking of target errors (Diedrichsen *et al*., 2005). Of particular interest is the finding that POS is a key player in PA process; according to a study by Luauté *et al*. (2009), POS activation is responsible for successful error correction during PA exposure, suggesting that it contributes to the strategic component of adaptation, which could mediate PA aftereffects on cognitive spatial representations of real-life environment. Furthermore, the fact that in our study POS area was activated especially in the right hemisphere could be interpreted as an enhanced representation of the left portion of space following rightward adaptation. Though in the present study, the analyses of naturalistic stimuli processing were limited to the rightward-VRPA group, future studies examining the effect of leftward-VRPA will determine whether the laterality of the neural aftereffects is dependent upon the directionality of the reference frame shift (i.e., by demonstrating symmetrical neural aftereffects), or it is generally lateralized to the right hemisphere.

A possible confounding factor that could influence our results is the fact that participants viewed the same exact set of videos twice, namely that there was a test-retest effect. However, data from the sham-VRPA group, who underwent exactly the same experimental paradigm, but without any adaptation shift, showed only highly inconsistent bilateral activity increases in the IPS area. Since the IPS area is highly activated both during reaching movements and during movie viewing (Culham *et al*., 2003; Goldberg *et al*., 2014; Nardo *et al*., 2016), one option is that this effect resonates activations of visuomotor practice during the sham training, as reflected also in a slight increase in visuomotor resting state connectivity.

Since participants were free to move their eyes during video presentation, another potential interpretation of our results is that the right POS enhancement after VRPA was due to differences in eye movement scanning patterns between pre- and post-VRPA training sessions (e.g., tendency to saccade more to the left after adaptation; as demonstrated for neglect patients in Serino *et al*., 2006). Unfortunately, we could not monitor eye movements in the scanner, so we cannot directly confirm, nor exclude this hypothesis. However, a recent study by Gilligan *et al*. (2019) found that PA had no effect on subsequent gaze directions in healthy individuals during passive gazing, but only when gazing involved active arm movements. Moreover, theoretically, only videos that presented bilateral salient objects actually lead to a competition between execution of left or right saccades (Nardo *et al*., 2016). But in our case, unilateral videos presentation naturally drew the gaze to the only salient moving stimulus, no matter whether it appeared on the right or on the left side of the screen. Therefore, the fact that the right hemisphere POS enhancement was evident for both left and right unilateral videos suggests that the neural aftereffects did not rely solely on differences in saccade directions. Instead, we suggest that the right POS upregulation might have stemmed from top-down influences from areas related to reference frame transformation and attention allocation, leading to an enhancement of left hemispace visual processing (Cohen & Andersen, 2002; Kim & Kastner, 2019).

#### Link between neural and behavioural aftereffects

A first indirect support for this hypothesis comes from the correlation between the behavioural aftereffects measured in the straight-ahead task and the upregulation of the right POS during naturalistic stimuli processing. Pointing straight ahead task is a common tool for assessing egocentric lateral spatial biases in both healthy individuals and neglect patients, representing proprioceptive-only effects (Bultitude *et al*., 2021), independent of any visual context (Karnath, 1994; Fleury *et al*., 2019). On the other hand, the movie viewing task was purely perceptual, and did not involve any manual or proprioceptive components. Thus, the fact that the magnitude of proprioceptive bias was proportional to the magnitude of right POS visual activations upregulation, suggests a common VRPA-induced over-representation of left space, manifested in both visual perception and proprioception. It is well accepted that PA induces a contralateral shift in reference frame (Redding & Wallace, 1996), which generalizes across different tasks (Michel, 2015). Thus, we propose that the same effect induced by VRPA manifests both for encoding of incoming stimuli and for movement execution. This change in reference frame might be mediated by top-down influences from higher order hetermodal regions (Cohen & Andersen, 2002), as tested via PPI analysis.

### 3. Naturalistic stimulus responses relate to connectivity modulation through DMN areas

PPI analysis demonstrated that following rightward VRPA, significant interactions emerged between right POS and areas in the DMN network during movies presentation, in areas that also showed VRPA-related changes in resting state connectivity. More specifically, right POS was found to interact with left IPL and bilateral STS regions (see *Figure 6*). Previous studies on PA-related brain activity found that activity in left IPL during visual attention task is modulated by PA in both healthy individuals and neglect patients (Crottaz-Herbette *et al*., 2014; Crottaz-Herbette *et al*., 2017a). Additionally, both IPL and STS change their restingstate connectivity following PA (Wilf *et al*., 2019), and have an important role in the representation of space. IPL is a key area for reference frames transformation (Cohen & Andersen, 2002). Its lesion results in severe spatial deficits in brain damaged patients (Corbetta *et al*., 2005), while its interference via TMS results in a lack of PA aftereffects in pointing straight ahead (Terruzzi *et al*., 2021). STS region is active during the last stages of PA (Luauté *et al*., 2009), and has been suggested to be the area integrating egocentric and object-centred reference frames (Karnath *et al*., 2001). Thus, these high-level IPL and STS regions are likely involved in mediating the shift in reference frames induced by sensorimotor adaption, which manifests multimodally in any spatial task, such as processing of naturalistic movies or pointing straight ahead. These internal biases are evident also in the spontaneous connectivity modulations, in the decoupling of DMN and attentional networks.

In addition, our results show that the connectivity changes bias hemispheric laterality depending on the direction of the adaptation. Therefore, a decrease in connectivity within the left hemisphere following rightward-VRPA might bias DMN connectivity in favor of the right hemisphere. This, in turn, can mediate enhanced activity in the right hemisphere in response to an incoming stimulus (*Figure 7*). This view is in line with a recent account of DMN function, namely its role in the integration of past information with current stimulus processing during naturalistic stimulation (Yeshurun *et al*., 2021). This function is supposed to be mediated by topographic connectivity between DMN and primary visual areas (Knapen, 2021), structuring the communication between distant brain regions. Our findings are in accordance with this model, with the addition of demonstrating that this long-range DMN-visual cortex organization adapts in cases of altered spatial representations, such as after VRPA.

**Figure 7.**
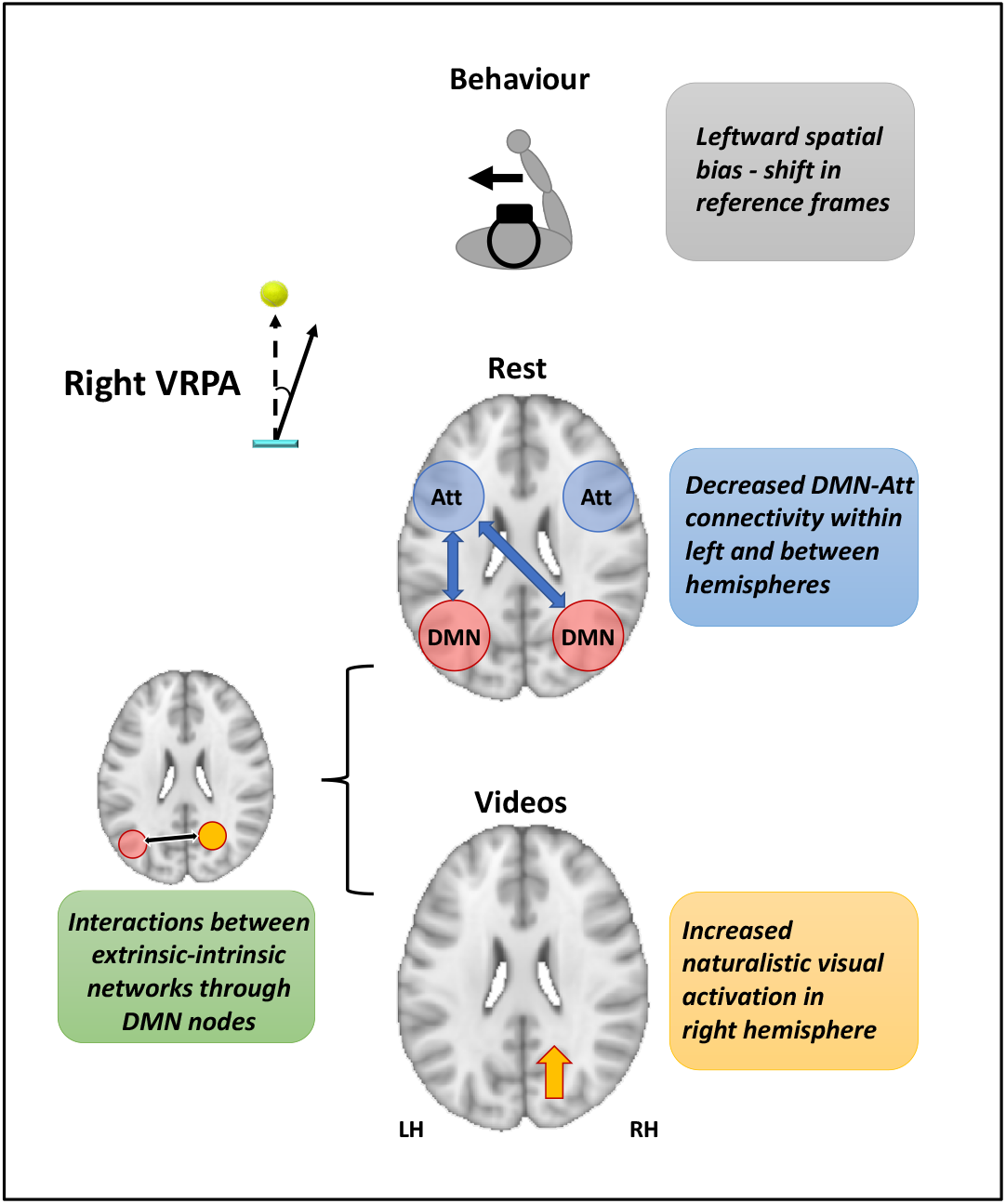
Schematic model for right-VRPA induced modulations. Bottom-up inputs during right VRPA training recalibrate spatial reference frames to the left side, and propagate to high level regions in the DMN and attentional networks (‘Att’). This, in turn, manifests in biases in resting state connectivity between DMN and Att, which mediates right hemisphere activity increase during naturalistic videos processing.

### 4. Potential implications for neglect rehabilitation

Long-range connectivity, mainly between regions of the DMN and the dorsal attentional system, is structurally altered in neglect syndrome, usually following a lesion in right frontoparietal regions, such as temporo-parietal junction (TPJ) (Corbetta *et al*., 2005; Corbetta & Shulman, 2011; Baldassarre *et al*., 2014). These unilateral lesions result in a general imbalance of functional connectivity between hemispheres (Lunven & Bartolomeo, 2017), that is proportional to the level of spatial impairment (Ptak *et al*., 2020). Accordingly, neglect recovery is associated to the re-emergence of decoupling between DMN and attentional networks within and across hemispheres (Ramsey *et al*., 2016). We propose that PA, and its current extension in VRPA, can modulate exactly these large-scale networks in favor of recreating connectivity changes similar to those evident during spontaneous recovery (see also Wilf *et al*., 2019). Importantly, the hemispheric laterality found in the connectivity modulations implies that this tool can be used, at least partially, to re-balance aberrant connectivity in neglect patients via compensatory mechanisms in the left DMN-attentional networks connectivity and the right visual cortex (Crottaz-Herbette *et al*., 2017a; Robineau *et al*., 2019). Unlike standard PA, our novel VRPA setup was designed to reproduce naturalistic ecological conditions – with the participant’s hand embodied as a virtual hand, and three-dimensional hand and target movements promoting natural behaviour (e.g., Carter *et al*., 2016). Hence, the VR setup have enabled us to tap into sensorimotor mechanisms that are more ecological than during standard PA (reaching to dynamic objects), thus potentially resulting in better transfer to real-life behaviour (Fortis *et al*., 2010; Champod *et al*., 2018). Though the adaptation shift was implemented using visuomotor rotation (cf., Krakauer, 2009; Gammeri *et al*., 2018), the fact that the feedback from the hand position was limited only to the last part of the movement (where the lateral shift was already close to its maximum size) promotes explicit adaptation mechanisms (Taylor *et al*., 2014), resembling the classic PA (Wilf *et al*., 2020).

In terms of visual processing, although the visual cortex is usually spared in patients with neglect without hemianopia, interferences in primary visual cortex activity were still found in certain cases (Vuilleumier *et al*., 2008). The right hemisphere enhancement we found following adaptation suggests that VRPA could potentially boost the damaged right visual cortex activation in neglect patients, through top-down modulations by fronto-parietal regions (Corbetta & Shulman, 2011). Naturalistic movies are a powerful tool to uncover pathologies in perception and biases in spatial orientation in the absence of an explicit task (cf., Machner *et al*., 2012; Nardo *et al*., 2019). Therefore, naturalistic stimuli can serve as a probe for daily-life perception and potential biases in spatial representations in neglect patients, and consequently track the potential re-balancing effect of VRPA rehabilitation training in these patients.

## Materials and Methods

### 1. Participants

The study comprised 45 healthy young adults, which were divided into three independent groups of participants: a first group of 16 participants performed rightward adaptation training (aged 23±3; 8 females), a second group of 14 participants performed sham training (aged 24±4; 7 females), and a third group of 15 participants performed leftward adaptation training (aged 23±4; 8 females). Required sample size was estimated based on our previous study that measured resting state modulations following standard PA (Wilf *et al*., 2019). All participants were right-handed with normal or corrected-to-normal vision and no neurological or psychiatric pathologies. Participants gave written informed consent according to procedures approved by the local Ethics Committee (CER-VD protocol no. 2017-01588). All participants were naïve about the purpose of the study, and had no prior visuomotor adaptation experience.

### 2. Procedures

#### 2.1. Experimental session overview

An experimental session lasted about 120 minutes, and included two phases of VR training interleaved with two identical phases of fMRI (pre-post adaptation).

##### Virtual Reality Device and setup

During the virtual reality training and tests, participants were seated on a chair in a fixed position, while the experiment was presented via a head mounted display (‘Oculus Rift’ Consumer Version 1, 1080×1200 resolution per eye, 110° Field of View, refresh rate of 90Hz). Participants were immersed in a unique 3D VR environment depicting a tennis field, which was developed using Unity^®^ Software (consumer version 2019.2.13), and used their right hand to control a virtual hand in the virtual world (see *Figure 1a*; Oculus Touch by Oculus).

The sequence of experiments was as follows (see *Figure 1b*):

###### Baseline VR phase

At the beginning of the experiment, participants performed 60 trials of baseline VR training outside the MRI without any visuomotor shift, during which they had to use the virtual hand to catch tennis balls that were thrown at them from the other side of the tennis field (see section ‘VRPA training’ for details). Immediately following the training, participants performed several open loop reaching tests, with no feedback regarding their hand position, in order to establish their baseline lateral spatial biases prior to adaptation. The aftereffect tests included reaching straight ahead with eyes closed, and open-loop pointing to left and right circles without visual feedback inside and outside VR (the open-loop results will not be covered in the current paper).

###### Pre-VRPA fMRI phase

The fMRI session started with an 8 min run of resting state with eyes closed, followed by a 7 min run of visual attention task (not covered in this paper), and a 9 min naturalistic stimulus run, in which participants freely viewed a series of short videoclips portraying everyday life scenes (videos taken from Nardo *et al*., 2016; Nardo *et al*., 2019; see Figure 1c).

###### VRPA phase

Participants went out of the MRI for an additional VR training phase, this time performing 160 trials of the tennis game with a 20° rotational shift between the virtual and their real hand, namely visuomotor adaptation (see section ‘VRPA training’ for details; no shift was introduced for the sham group), followed by identical open loop tests to probe adaptation-induced behavioural aftereffects.

###### Post-VR-adaptation fMRI phase

Immediately following the VR session, participants went back in the MRI (they had to cross their hands and were not allowed to use them during the transition in order to avoid potential de-adaptation). Participants repeated the same fMRI experimental sequence as in the first fMRI session - i.e., rest with eyes closed, visual attention task, and naturalistic viewing task.

###### Final open-loop tests

Additional repetition of the open loop tests was performed after the second fMRI phase, to assess the residual behavioural aftereffects also at the end of the fMRI session, i.e., about 40 min past the adaptation phase.

The experimental sequence described above applies to the right-VRPA and sham-VRPA groups. However, the left VRPA group performed a different paradigm with only resting state and different visual tasks during the fMRI sessions.

#### 2.2. VRPA training

During the VR training phases, participants were immersed in a VR environment of a tennis-field using a natural-looking virtual hand, and were instructed to catch tennis balls that were thrown at them from the far end of the field. Each trial began when participants placed their hand in the ‘origin position’ located a few centimeters in front of the midline of their body, at position [0, 0] on the XZ plane of the virtual environment. To guide participants back to the correct origin position, the area around it was marked by a semi-transparent turquoise semicircle (30 cm radius). The following trial was launched only when participant’s hand was placed back in the origin position. Importantly, the participants’ virtual hand was invisible while the hand was within the radius of the round table (~30 cm from the origin position), and became visible only in the last segment of the reaching movement trajectory, namely when the hand was already close to the target. This partial visual feedback of hand trajectory enables both strategic and online mechanisms to take place during adaptation (Facchin *et al*., 2020; Wilf *et al*., 2020). In each trial, the tennis ball approached the participant at a constant speed randomly selected within the range of comfortable ball speeds. The balls were thrown from one of four initial positions - either the far left position/the far right position, above/below participant’s arm height (initial positions ranged from −2m to 2m on the x axis, and 0.2m to 1.8m on the y axis, equidistant 4m from the origin position), and moved towards one of 24 possible terminal positions equally spread around the midline, resulting in an equal number of balls moving from left to left, left to right, right to right, and right to left hemifields (*Figure 1a*; terminal positions ranged from −0.4m to 0.4m on the x axis, and from 1m to 1.2m on the y axis, equidistant 0.45 m from the origin position). The order of trials was randomized in each session. When catching the ball, participants received multisensory positive feedback: a wind-chime auditory sound, a light vibration tactile feedback of the controller, and a bright glitter visual effect. Participants were able to catch the balls also before or after they reached the intended terminal positions, but in case a ball was not caught, it either continued to move and disappeared behind the participant, or it ‘blew up’ before it reached the turquois area surrounding the subject. The radius for catching the targets was limited to a minimum of 30 cm in order to force the participants to perform large and quick reaching movements that enable proper adaptation, and to prevent the ball from arriving too close to the participant’s body. During the VRPA session, a rightward/leftward 20° rotational shift was induced between the real hand and the virtual hand (see *Figure 1a*) on the 11^th^ trial (similarly to Wilf *et al*., 2020)). During baseline and sham-VRPA sessions, the virtual hand and the real hand were always aligned.

#### 2.3. Subjective midline test

Participants were instructed to close their eyes and reach straight ahead in front of the midline of their body. Once they reached their subjective midline position, they pressed the trigger button of the VR controller with their index finger, and returned their hand in the origin position near their body. The pointing error was calculated as the angular deviation from the true midline (referred to as 0 on the x axis, directly in front of the origin position). This procedure was repeated 5 times, and the five movements were averaged to establish the participant’s pointing error in each of the phases of the experiment (see *Figure 1b*): pre (immediately following the VR-baseline session), post (immediately following the VRPA session), and end (following the second fMRI session, ~40 minutes following adaptation).

An additional open loop aftereffect test was performed outside the VR environment, where participants had to look at either left or right targets and then reach them with their eyes closed (see Crottaz-Herbette *et al*., 2014 for standard open loop test procedure). The angular target error at each experimental phase was calculated as in the subjective midline task.

A 2-way repeated measures ANOVA was performed for each of the aftereffect tests separately using the JASP software version 0.11.1, with within-subject factor phase (pre/post/end) and between subjects factor group (right VRPA/sham VRPA), followed by post-hoc tests.

### 3. Imaging Setup and fMRI Data analysis

MRI data of right-VRPA and sham-VRPA groups was collected with a 3T Siemens Magnetom Prisma scanner with a 64-channel head-coil, located at the Lemanic Biomedical Imaging Center (CIBM), and data of group left-VRPA was collected at the Laboratory for Research in Neuroimaging (LREN) in the Centre Hospitalier Universitaire Vaudois (CHUV), Lausanne. Functional MR images were acquired with a multiband-2 echo planar imaging gradient echo sequence (repetition time 2 s; flip angle 90°; echo time 30 ms; number of slices 66; voxel size 2×2×2.04mm; 10% gap). The 66 slices were acquired in a sequential ascending order and covered the whole head volume in the AC-PC plane. A high-resolution T1-weighted 3D gradient-echo sequence was acquired for each participant (right VRPA group: MP2RAGE as described in (Marques *et al*., 2010); sham and left VRPA groups: MPRAGE, voxel size 1×1×1 mm). To prevent head movements in the coil, padding was placed around the participant’s head.

#### 3.1. Experiment 1: fMRI Data analysis resting state data

##### 3.1.1. MRI data preprocessing

Data were processed using FSL 5.0.10 (www.fmrib.ox.ac.uk/fsl) and in-house Matlab code (Mathworks, Natick, MA, USA). Functional data of all groups was analyzed with an identical pipeline, while for right VRPA group, due to the use of MP2RAGE, an additional step was taken in the analysis of the anatomical scans in order to unite the sparse MP2RAGE images into a single anatomical file (brightness threshold 90, and multiply INV2 and UNI for noise removal; Marques *et al*., 2010).

Functional data were analyzed using FMRIB’s expert analysis tool (FEAT, version 6). The following pre-statistics processing was applied to the data of each participant: motion correction using FMRIB’s Linear Image Registration Tool (MCFLIRT) (Jenkinson et al., 2002); brain extraction using BET (Smith et al. 2004). For right VRPA group, the BET was performed on the INV2 contrast file for better brain extraction (Choi *et al*., 2019); high-pass temporal filtering with a cut-off frequency of 0.01 Hz; removal of the first 2 volumes from each functional run, and 5 mm Gaussian spatial smoothing. Functional images were aligned with high-resolution anatomical volumes initially using linear registration (FLIRT), then optimized using Boundary-Based Registration. Structural images were then transformed into standard MNI space using non-linear registration tool (FNIRT), and the resulting warp parameters were applied to the functional images as well. All the functional images were resampled to 2×2×2 mm^3^ standard space, and therefore all further analyses were performed in standard MNI space.

Only for resting-state data, additional denoising steps were performed: A scrubbing procedure was applied for censoring motion-contaminated frames using the framewise displacement (FD) and the DVARS measures (Power *et al*., 2014). Then regressing out of signals from the white matter and ventricles was performed as follows: The white matter and ventricles of each participant were automatically defined using FSL’s FAST (Zhang *et al*., 2001), and eroded to avoid boundaries between tissues (Hahamy et al., 2015). The non-neuronal contributions to the BOLD signal were removed by linear regression of motion parameters, ventricle and white matter timecourses for each participant (Fox *et al*., 2009), while no global signal regression was performed (Weissenbacher *et al*., 2009; Hahamy *et al*., 2014; Murphy & Fox, 2017). Two participants from the right VRPA group were removed from analysis due to prolonged contamination by excessive head motion (> 4mm), resulting in 14 participants for the resting state right-VRPA group, and 14 participants for the resting state sham group.

##### 3.1.2. Atlas-based segmentation and network labeling

Cortical regions of interest (ROIs) for resting-state connectivity analysis were defined based on the Harvard-Oxford probabilistic atlas (as implemented in FSL software; Desikan *et al*., 2006). The definition of regions was performed for each participant as following: First, the atlas label mask was multiplied by the individual subject’s gray matter mask (as defined by FSL’s FAST) to exclude voxels outside the brain. Then, to enable comparison between left and right hemispheres, the 48 atlas-defined regions were each separated to two distinct regions in the left and right hemisphere according to x-coordinates of the voxels comprising each region, resulting in 48 pairs of homologue regions symmetrical across the two hemispheres (see *Figure 3a*). In order to sort the regions in a functionally-relevant manner, each region was assigned to one of seven primary functional connectivity resting state networks based on the maximum overlap of the region with the network MNI cortical parcellation (Yeo *et al*., 2011): *visual, somatomotor, dorsal attention network (DAN), ventral attention network (VAN), limbic system, frontoparietal network (FPN), and default mode* network (DMN). Within each network, regions were sorted according to functional subsystems, and arranged from posterior to anterior and from dorsal to ventral (see *Figure 3a* and *Supplementary Table 1* for full sorted list of regions).

##### 3.1.3. Functional connectivity matrices

A connectivity matrix was calculated for each resting state run in each participant as follows: the mean timecourse of each region was extracted, and pairwise Pearson correlation was calculated for each pair out of the atlas regions, resulting in a 96×96 correlation matrix (48 left hemisphere and 48 right hemisphere regions). The order of regions in the matrix remained identical in the left and right hemisphere regions. This resulted in a symmetrical matrix, with upper left quadrant depicting within-hemisphere connectivity in the left hemisphere (‘LL’), lower right quadrant depicting within-hemisphere connectivity in the right hemisphere (‘RR’), and lower left quadrant depicting connectivity between left and right hemisphere (‘LR’, the quadrant diagonal represents homotopic connectivity between homologue regions of the two hemispheres; see scheme in *Figure 4c*).

The single-subject connectivity matrices were Fisher z-transformed and averaged to generate the mean ‘pre’ and ‘post’ resting state connectivity matrices. Each individual participant’s ‘post’ and ‘pre’ matrices were subtracted, and the modulation matrices were averaged to visualize the average difference in connectivity between ‘pre’ and ‘post’.

##### 3.1.4. Gaussian mixture model analysis

For assessing the overall distribution of correlation coefficients of the ‘pre’ and ‘post’ single-subject connectivity matrices, a Gaussian mixture modeling (GMM) approach was implemented (Tyszka *et al*., 2014). First, to avoid redundancy, only the values below the main diagonal were taken for further analysis (see schema in *Figure 4c*). Then the distribution of correlation coefficients in each matrix was plotted, and fitted with a combination of 2 Gaussian components (see example in *Figure 4a*; as implemented in Matlab Statistical Toolbox with 3000 iterations and 100 replications). Then for each matrix the μ (mean of the Gaussian) accounting for the highest proportion of the data variability was taken as the representative value of the matrix distribution, resulting in two μ values for each subject that were taken for further statistical analysis (‘pre’ and ‘post’). A 2-way mixed-effect repeated-measures ANOVA (rmANOVA) was then performed using JASP with factors ‘phase’ (pre/post; within-subject factor) and group (right-VRPA/sham-VRPA; between-subject factor).

To quantify the change in connectivity within each hemisphere and between hemispheres, an analysis based on the same principle was performed separately for each quadrant of the whole connectivity matrix, but this time plotting the distribution of the *difference* matrices (post-pre). The matrices were divided to three quadrants – i.e., correlation values depicting left-left connections (LL), right-right connections (RR), and interhemispheric left-right connections (RL; see *Figure 4c; right*). Each Gaussian fit was performed separately for each quadrant, resulting in 3 μ values for each subject denoting the difference between ‘pre’ and ‘post’ (for LL, RR, and RL). For the LL and RR submatrices, only values below the diagonal were considered, but since the interhemispheric submatrix RL is not entirely symmetrical, all its values were included in the distribution and not only values below the subdiagonal (note that homotopic connections along the submatrix diagonal were also included, since they are not equal to 1 and therefore not necessarily cancel out when subtracting ‘pre’ and ‘post’ matrices). The resulting μ parameters were taken to statistical analysis of a 2-way mixed-effect rmANOVA with factors ‘quadrant’ (LL/RR/RL; within-subject) and group (left-VRPA/right-VRPA/sham-VRPA; between-subject).

#### 3.2. Experiment 2: free viewing of naturalistic videos

##### 3.2.1. Experimental Design and stimuli

During the pre- and post-fMRI sessions, participants were presented with videos portraying everyday life scenes, and were instructed to freely view the videos without any explicit task. We used a well-validated set of visual-only short videoclips (1.5 sec duration), in which salient objects appeared either on the right side, left side, or both sides of the screen (Nardo *et al*., 2016; Nardo *et al*., 2019). Out of the original stimulus set, 80 videos were selected, divided into groups of 20 videos from four different categories according to previously established saliency maps (Nardo *et al*., 2016): left-lateralized, right-lateralized, bilateral left salient, bilateral right salient; see Figure 1b), while ensuring a balanced distribution of objects representing different visual categories (faces, cars, body-parts etc..). Each run began with one 3 sec video of visual moving texture, to eliminate the transient response to a novel visual stimulation, which might artificially increase the response to the first trial. The order of videos was pseudorandomized and videos were interleaved with periods of fixation lasting between 4.5-8.5 sec (event timings were determined using “optseq” tool (Dale *et al*., 1999)). The content of left-salient and right-salient videos was counterbalanced by presenting a horizontally-flipped version of the videos to half of the participants (however, note that each participant viewed an identical version of the videos in the pre- and post-sessions).

##### 3.2.2. MRI data preprocessing

Preprocessing was similar to the one described in section 3.1.1 for resting state data, with the exception that projecting out and scrubbing were not perform on task-based data, but instead noise factors were controlled through the event-based model. One participant from the right VRPA group was removed from analysis due to prolonged contamination by excessive head motion (> 4mm), resulting in 15 participants for the videos right VRPA group, and 14 participants for the videos sham group.

##### 3.2.3. Multisubject general linear model (GLM) analysis

To create task-based statistical parametric maps, a whole brain GLM analysis was applied for each subject using FEAT, modeling each video category with a corresponding regressor (convolved with double-gamma hemodynamic response function). The six motion parameters and their derivatives were used as nuisance regressors. In addition, motion-contaminated timepoints were modeled as nuisance regressors using motion outlier detection algorithm implemented in FSL.

This analysis yielded five statistical maps of interest: response to all videos, to left-lateralized videos, right-lateralized videos, Left-salient bilateral videos, and right-salient bilateral videos. Since each participants underwent two identical runs of video presentation, the single-subject analysis resulted in two maps for each participant in each condition (‘pre’ and ‘post’). A group analysis of ‘pre’ session and ‘post’ session separately was carried out using FMRIB’s Local Analysis of Mixed Effects (FLAME1). Z statistic images were thresholded using clusters determined by Z > 2.6, and a family-wise-error corrected cluster significance threshold of p<0.05 was applied to the suprathreshold clusters.

To compare between ‘pre’ and ‘post’ sessions, a within-subject analysis of statistical maps was carried out using a paired two-group difference design in FMRIB’s Mixed Effect Ordinary Least Squares Estimation, cluster thresholded Z > 2.3 and p < 0.05 (results were verified and replicated also using SPM paired t-test GLM with threshold 0.005 FDR correction).

Each of the four experimental condition was examined separately (left/right/unilateral/bilateral videos), as well as joined together to see general responses to naturalistic videos regardless of their spatial layout.

##### 3.2.4. Correlation between brain and behaviour

The change in cortical activity during movie presentation (‘all videos’ condition) was compared to the change in subjective midline open loop pointing in our group of participants. To that end, a region of interest (ROI) was defined according to the group map generated by contrasting ‘pre’ and ‘post’ responses to all videos (separately for the right VR-adaptation group and for the sham group). The beta value of this ROI from each individual participant was extracted in the ‘pre’ and ‘post’ runs. Then the difference in beta value (‘post’-‘pre’) was compared to the difference in open loop pointing error (deviation from true midline) by means of Pearson correlation using JASP software for statistical analysis.

##### 3.2.5. Psychophysiological Interaction (PPI) analysis

PPI analysis was carried out using FSL FEAT, to probe possible interactions between the selected ROI and the rest of the brain, related to videos presentation (right POS ROI was taken for right VR-adaptation group, bilateral IPS ROI for sham group). To that end, the activity timecourse of the ROI was extracted in each participant and each run (‘pre’ and ‘post’), and its interaction with an ‘all videos’ regressor was assessed. Then a group GLM analysis was performed on the ‘pre’ and ‘post’ sessions with FLAME1 (Z > 2.3, FWER p<0.05 cluster correction).

## Acknowledgements

This work was supported by the European Union’s Horizon 2020 research and innovation programme under the Marie Sklodowska-Curie grant agreement No. 789548 and the Swiss government excellence scholarship to M.W, and the Swiss National Science Foundation grant (grant PP00P3_163951/1) to A.S. The authors wish to thank all the participants who took part in the study. The authors thank Antoine Lutti, Giulia Di Domenicantonio, Christine Kieffer, Petr Grivaz, and Giulio Mastria for their help in data acquisition, and Dimitri Van de Ville for helpful comments and feedback on data analysis.

## Notes

### Competing Interest Statement

The authors have declared no competing interest.

