## Supplementary Figures and Table for "Virtual reality-based sensorimotor adaptation shapes subsequent spontaneous and naturalistic stimulus-driven brain activity"

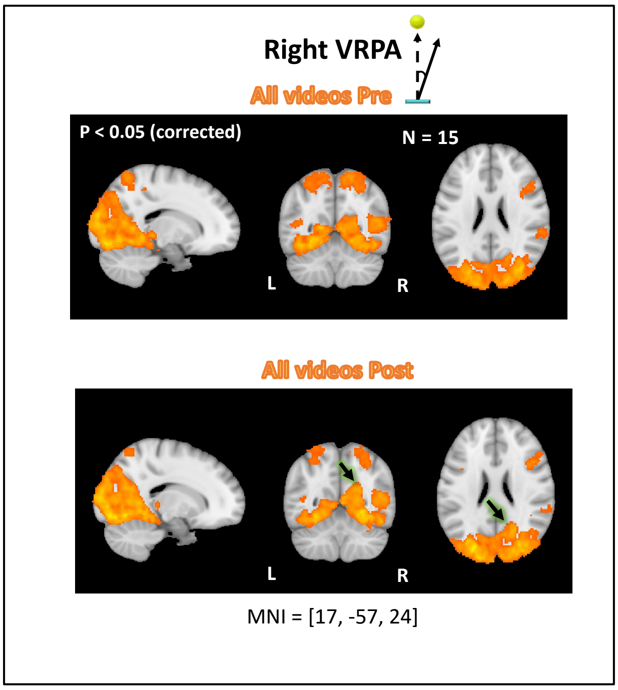


***Supplementary Figure 1. Activation in response to free viewing of videos before and after right VRPA****. Activation maps for all video types versus rest in the right VRPA group (N=15), before (PRE) and after (POST) VRPA training. Green-black arrow denotes the area in which activation was enhanced following right VRPA.*

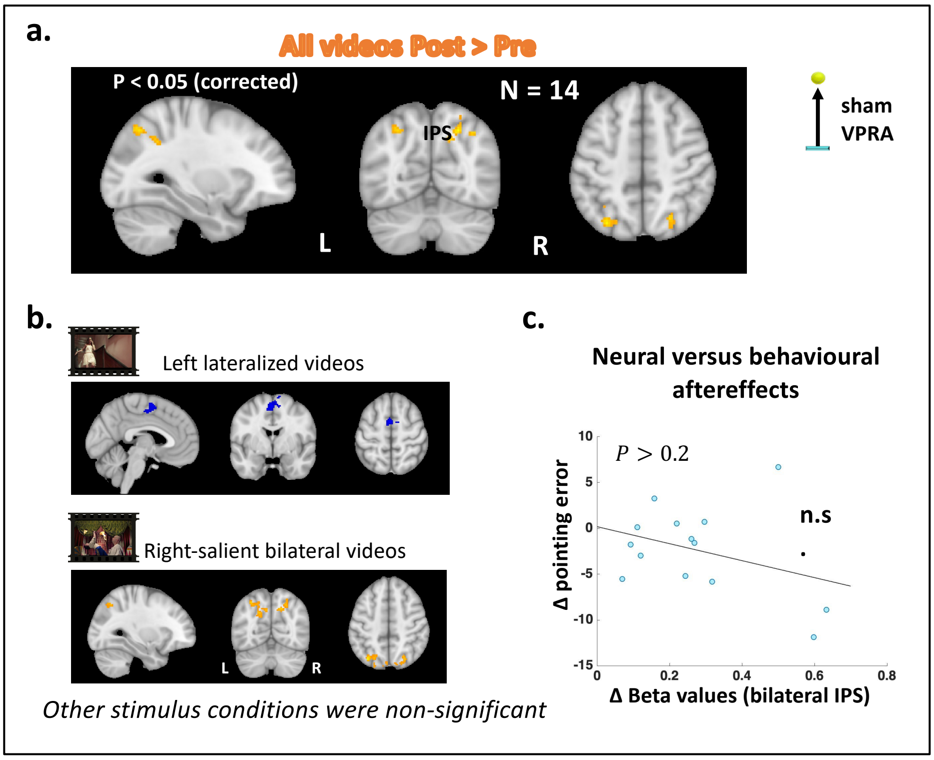


***Supplementary Figure 2. Sham-VRPA induced inconsistent modulations of brain activity in response to videos****. (a) Direct comparison between responses to all video types post- and pre- sham-VRPA showed enhanced bilateral IPS activation. (b) Response modulation in each video condition. Only right-salient bilateral videos showed enhancements in activation, while the other video condition showed no significant modulation or even an activation decrease (blue shades in 'left-lateralized' map). (c) correlation between bilateral IPS and changes in pointing error showed no correlation (same specifications as in Figure 5). IPS = Intraparietal Sulcus.*

*Supplementary Table 1:*

| Matrix serial number | H-O atlas number | Harvard-Oxford atlas label | Yeo network label | hemisphere |
|---|---|---|---|---|
| 1 | 48 | occipital pole | visual | L |
| 2 | 47 | supracalcarine | visual | L |
| 3 | 24 | intracalcarine | visual | L |
| 4 | 36 | lingual gyrus | visual | L |
| 5 | 23 | LOC | visual | L |
| 6 | 40 | fusiform gyrus | visual | L |
| 7 | 39 | temporal occipital fusiform | visual | L |
| 8 | 32 | cuneus | visual | L |
| 9 | 17 | postcentral gyrus | somatomotor | L |
| 10 | 7 | precentral gyrus | somatomotor | L |
| 11 | 26 | SMA | somatomotor | L |
| 12 | 45 | heschl's gyrus | somatomotor | L |
| 13 | 46 | planum temporale | somatomotor | L |
| 14 | 44 | planum polare | somatomotor | L |
| 15 | 42 | central operculus | somatomotor | L |
| 16 | 9 | anterior STG | somatomotor | L |
| 17 | 10 | posterior STG | somatomotor | L |
| 18 | 16 | Temporo-occipital ITG | DAN | L |
| 19 | 19 | anterior SMG | DAN | L |
| 20 | 18 | SPL | DAN | L |
| 21 | 13 | Temporo-occipital MTG | DAN | L |
| 22 | 2 | insula | VAN | L |
| 23 | 43 | parietal operculum | VAN | L |
| 24 | 20 | posterior SMG | VAN | L |
| 25 | 15 | posterior ITG | limbic | L |
| 26 | 14 | anterior ITG | limbic | L |
| 27 | 34 | anterior PHG | limbic | L |
| 28 | 37 | anterior temporal fusiform | limbic | L |
| 29 | 38 | posterior temporal fusiform | limbic | L |
| 30 | 8 | temporal pole | limbic | L |
| 31 | 27 | subcallosar cortex | limbic | L |
| 32 | 33 | orbital | limbic | L |
| 33 | 41 | frontal operculum | FPN | L |
| 34 | 5 | IFG triangularis | FPN | L |
| 35 | 6 | IFG opercularis | FPN | L |
| 36 | 29 | anterior cingulate | FPN | L |
| 37 | 28 | paracingulate | FPN | L |
| 38 | 1 | frontal pole | FPN | L |
| 39 | 12 | posterior MTG | DMN | L |
| 40 | 11 | anterior MTG | DMN | L |

| | | | | |
|---|---|---|---|---|
| 41 | 31 | precuneus | DMN | L |
| 42 | 22 | IPL | DMN | L |
| 43 | 21 | angular gyrus | DMN | L |
| 44 | 30 | posterior cingulate | DMN | L |
| 45 | 25 | frontal medial cortex | DMN | L |
| 46 | 4 | MFG | DMN | L |
| 47 | 3 | SFG | DMN | L |
| 48 | 35 | posterior PHG | DMN | L |
| 49 | 48 | occipital pole | visual | R |
| 50 | 47 | supracalcarine | visual | R |
| 51 | 24 | intracalcarine | visual | R |
| 52 | 36 | lingual gyrus | visual | R |
| 53 | 23 | LOC | visual | R |
| 54 | 40 | fusiform gyrus | visual | R |
| 55 | 39 | temporal occipital fusiform | visual | R |
| 56 | 32 | cuneus | visual | R |
| 57 | 17 | postcentral gyrus | somatomotor | R |
| 58 | 7 | precentral gyrus | somatomotor | R |
| 59 | 26 | SMA | somatomotor | R |
| 60 | 45 | heschl's gyrus | somatomotor | R |
| 61 | 46 | planum temporale | somatomotor | R |
| 62 | 44 | planum polare | somatomotor | R |
| 63 | 42 | central operculus | somatomotor | R |
| 64 | 9 | anterior STG | somatomotor | R |
| 65 | 10 | posterior STG | somatomotor | R |
| 66 | 16 | Temporo-occipital ITG | DAN | R |
| 67 | 19 | anterior SMG | DAN | R |
| 68 | 18 | SPL | DAN | R |
| 69 | 13 | Temporo-occipital MTG | DAN | R |
| 70 | 2 | insula | VAN | R |
| 71 | 43 | parietal operculum | VAN | R |
| 72 | 20 | posterior SMG | VAN | R |
| 73 | 15 | posterior ITG | limbic | R |
| 74 | 14 | anterior ITG | limbic | R |
| 75 | 34 | anterior PHG | limbic | R |
| 76 | 37 | anterior temporal fusiform | limbic | R |
| 77 | 38 | posterior temporal fusiform | limbic | R |
| 78 | 8 | temporal pole | limbic | R |
| 79 | 27 | subcallosar cortex | limbic | R |
| 80 | 33 | orbital | limbic | R |
| 81 | 41 | frontal operculum | FPN | R |
| 82 | 5 | IFG triangularis | FPN | R |
| 83 | 6 | IFG opercularis | FPN | R |

| | | | | |
|---|---|---|---|---|
| 84 | 29 | anterior cingulate | FPN | R |
| 85 | 28 | paracingulate | FPN | R |
| 86 | 1 | frontal pole | FPN | R |
| 87 | 12 | posterior MTG | DMN | R |
| 88 | 11 | anterior MTG | DMN | R |
| 89 | 31 | precuneus | DMN | R |
| 90 | 22 | IPL | DMN | R |
| 91 | 21 | angular gyrus | DMN | R |
| 92 | 30 | posterior cingulate | DMN | R |
| 93 | 25 | frontal medial cortex | DMN | R |
| 94 | 4 | MFG | DMN | R |
| 95 | 3 | SFG | DMN | R |
| 96 | 35 | posterior PHG | DMN | R |
